# A Connectome-Constrained Jansen–Rit Framework for Inferring Cortical Gain Control and Ensemble Stability

**DOI:** 10.64898/2026.01.28.702100

**Authors:** Andreea O. Diaconescu, Zheng Wang, John D. Griffiths

**Affiliations:** Krembil Centre for Neuroinformatics, Centre for Addiction and Mental Health (CAMH), Toronto, Ontario, Canada; Institute of Medical Sciences, University of Toronto, Toronto, Ontario, Canada; Department of Psychiatry, University of Toronto, Toronto, Ontario, Canada

## Abstract

Understanding how local circuit dynamics give rise to large-scale stability and instability of brain activity is a central challenge in computational neuroscience, with direct relevance for disorders characterized by disrupted excitatory–inhibitory balance, including schizophrenia spectrum disorder (SSD). Here, we introduce a principled methodology for recovering local neural parameters and low-dimensional dynamical biomarkers from a connectome-constrained Jansen–Rit (JR) neural mass model using variational free-energy inversion and sliding-window analysis. Each cortical region is modeled as a canonical excitatory–inhibitory microcircuit embedded within a whole-brain network whose long-range interactions are factorized into pyramidal–pyramidal, pyramidal– excitatory, and pyramidal–inhibitory subnetworks. Across 80 independent simulations, the inversion framework reliably recovered both microcircuit parameters and emergent biomarkers derived from neural states, including the mean–variance slope (*β*_1_), its spatial variability, and the lag-1 autocorrelation (*ρ*_1_). These quantities capture complementary aspects of cortical ensemble dynamics—gain sensitivity, regional heterogeneity, and temporal persistence associated with proximity to criticality—and were consistently estimated with minimal bias and high reliability. The recovered slope hierarchy 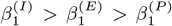 revealed an interpretable gain-control architecture in which inhibitory channels regulate damping, excitatory channels gate resonance, and pyramidal populations integrate network drive into stable output. Together, these results demonstrate that the JR model provides a tractable and biophysically grounded framework for linking synaptic parameters, network structure, and ensemble-level stability. Although motivated by questions surrounding psychosis risk and SSD, the proposed approach is general and establishes a foundation for future applications in model-based inference, network control, and adaptive neuromodulation.

## 1 Introduction

A central objective of computational neuroscience is to explain how alterations in neural circuit dynamics give rise to vulnerability for neuropsychiatric disorders. In the context of early psychosis and psychosis risk, converging evidence suggests that symptoms emerge not from isolated regional deficits, but from systematic disruptions in excitatory–inhibitory (E/I) regulation and the coordination of distributed neural ensembles Krystal et al. (2017); Uhlhaas and Singer (2010); Friston (2005); Adams et al. (2013). These disturbances are detectable in electrophysiological signals such as event-related potentials and oscillatory synchrony, and are hypothesized to reflect maladaptive shifts in circuit gain, temporal precision, and dynamical stability Uhlhaas et al. (2006, 2010); Taylor and Tso (2015); Gonzalez-Burgos and Lewis (2008). Despite substantial empirical progress, a key open challenge is to develop computational models that link such population-level observations to interpretable circuit mechanisms in a manner that is reproducible, identifiable, and scalable.

Neural ensemble theories provide a unifying framework for this challenge. In this view, cognitive and perceptual processes arise from the transient formation and stabilization of coordinated assemblies spanning multiple spatial and temporal scales Krystal et al. (2017). Psychosis risk is hypothesized to involve impaired formation, excessive instability, or inappropriate synchronization of these ensembles, leading to degraded contextual integration and aberrant inference Lisman et al. (2008); Rolls et al. (2008); Adams et al. (2013).

At the microcircuit level, altered glutamatergic and GABAergic transmission can shift cortical columns toward hyperexcitable or weakly damped regimes, compromising the balance between amplification and suppression required for stable ensemble dynamics Lewis et al. (2005); Gonzalez-Burgos et al. (2010); Bastos et al. (2012). At larger scales, these local perturbations propagate through anatomical connectivity, producing abnormal oscillatory coupling and reduced signal-to-noise ratio across networks Uhlhaas et al. (2009); Friston (2008).

From a dynamical-systems perspective, these effects can be interpreted as maladaptive regulation of operating points relative to critical manifolds. Cortical networks are thought to operate near criticality, where sensitivity, flexibility, and information transmission are maximized while stability is maintained Deco et al. (2011); Breakspear (2017). Systematic deviations from this regime—either toward excessive instability or overdamped dynamics—are predicted to impair ensemble coordination and may constitute a mechanistic signature of psychosis vulnerability Stephan et al. (2006); Heitmann and Breakspear (2018). Crucially, such deviations may be expressed not as large parameter changes but as subtle shifts in low-dimensional dynamical statistics, including gain sensitivity, variability, and temporal correlations.

To formalize these ideas, generative neural mass models provide a principled computational bridge between synaptic-level parameters and macroscopic neural signals measured with EEG or fMRI Jansen and Rit (1995); David and Friston (2003a); Moran et al. (2013a); Deco et al. (2011). Among these, the Jansen–Rit (JR) model offers a compact and biophysically interpretable description of E/I interactions within a cortical column Jansen and Rit (1995); David and Friston (2003b); Heitmann et al. (2017). When embedded within whole-brain networks, JR-based models have been used to study oscillations, metastability, and state transitions emerging from structured connectivity Breakspear (2017); Deco and Kringelbach (2017); Cabral et al. (2017); Breakspear et al. (2021). However, most applications either fix parameters heuristically or focus on steady-state behavior, limiting their utility for identifying dynamic biomarkers linked to ensemble stability and disease-relevant mechanisms.

Recent advances in differentiable simulators and variational inference enable model inversion frameworks that recover latent states and parameters directly from neural time series using free-energy optimization Deco and Kringelbach (2017); Nozari et al. (2020). These methods open the possibility of extracting low-dimensional, mechanistically grounded biomarkers of dynamical stability—such as gain sensitivity and critical slowing-down—from generative models, while retaining explicit links to underlying circuit parameters. Such an approach is well aligned with the goals of computational biology: to derive explanatory models that are both biologically interpretable and quantitatively testable.

Here, we develop a connectome-constrained Jansen–Rit modeling framework with variational Bayesian inversion to study how local circuit parameters give rise to emergent ensemble-level stability signatures. Long-range interactions are factorized into three functional subnetworks—pyramidal-to-pyramidal (P→P), pyramidal-to-excitatory (P→E), and pyramidal-to-inhibitory (P→I)—corresponding to distinct pathways that regulate baseline excitability, resonance gain, and damping Bastos et al. (2012). Within this framework, ensemble dynamics arise from the interaction between local microcircuit non-linearities and structured network input.

Using repeated model re-fitting, we assess the identifiability and reliability of three low-dimensional biomarkers derived from simulated neural states: the mean–variance slope (*β*_1_), its spatial variability, and the lag-1 autocorrelation (*ρ*_1_). These measures respectively index gain sensitivity, spatial heterogeneity, and temporal persistence associated with proximity to criticality, providing a compact description of ensemble organization that is directly linked to model parameters. We show that these biomarkers are robustly recovered across repeated inversions and reproduce a consistent hierarchy 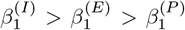 reflecting a gain-control architecture in which inhibitory channels operate closest to the critical boundary, excitatory channels gate resonance, and pyramidal populations integrate network drive into stable output.

By grounding neural ensemble theories of psychosis risk in a tractable biophysical model with reproducible inversion and interpretable dynamical readouts, this work contributes a general computational framework for linking E/I regulation, criticality, and ensemble stability. Although motivated by early psychosis, the modeling approach and biomarkers introduced here are broadly applicable to studies of large-scale brain dynamics and neuropsychiatric disorders characterized by altered circuit gain and stability.

## 2 Methods

### 2.1 Local JR Microcircuit (Cortical Only)

Each cortical parcel is modeled as a Jansen–Rit (JR) unit comprising three interacting populations: pyramidal cells (*P*), excitatory interneurons (*E*), and inhibitory interneurons (*I*). Presynaptic firing activity *v*(*t*) is transformed into postsynaptic potentials (PSPs) via second-order linear filters with excitatory and inhibitory time constants:

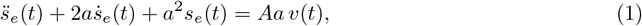

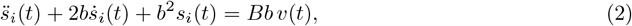

where *A* and *B* are excitatory and inhibitory synaptic gains, and *a* and *b* (s^*−*1^) are the corresponding inverse time constants.

Population firing rates are generated from mean membrane potentials using a static sigmoid:

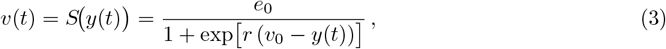

with maximal firing rate *e*_0_, half-activation potential *v*_0_, and slope *r*.

Within each parcel, local coupling is parameterized by four gains *C*_1_–*C*_4_, corresponding to the directed connections *P*→ *E* (*C*_1_), *E*→ *P* (*C*_2_), *P* →*I* (*C*_3_), and *I* →*P* (*C*_4_). Let *y*_*P*_, *y*_*E*_, and *y*_*I*_ denote the mean membrane potentials of the three populations. Their effective inputs are

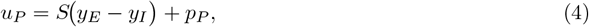

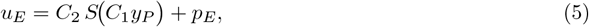

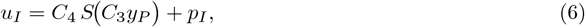

where *p*_*P*_, *p*_*E*_, and *p*_*I*_ represent external (tonic or stochastic) drives to the respective populations.

The population states then evolve according to JR-form second-order dynamics:

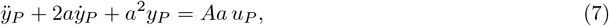

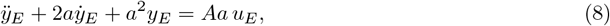

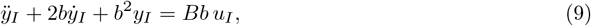

where long-range network inputs enter through the external drive terms as described in Sec. 2.2.

### 2.2 Connectome-Constrained Three-Network Factorization

In the Jansen–Rit (JR) neural mass model, each cortical region is represented by a three-population microcircuit (pyramidal projection neurons, excitatory feedback, and inhibitory feedback). In the original coupled-column formulation, inter-column coupling is implemented by adding a delayed, gain-scaled contribution from the sending column into the same afferent input channel as the external drive *p*(*t*), i.e. into the excitatory-loop equation that generates the net postsynaptic drive to pyramidal neurons (Jansen and Rit, 1995). This is the same strategy adopted in the connectome-embedded JR model of (Momi et al., 2023), where the long-range term conn_*j*_(*t*) is computed from delayed, weighted activity in other nodes and *enters the excitatory population only*.

Motivated by this canonical JR coupling location, we define an inter-regional coupling architecture with up to three extrinsic channels, all driven by the sending region’s pyramidal output, but targeting different subpopulations in the receiving region:

- **P**→ **E (canonical):** long-range excitation injected into the JR afferent input channel of the excitatory subpopulation (the same channel as *p*(*t*));
- **P**→ **I (extension):** long-range excitation targeting inhibitory feedback, modelling long-range recruitment of local inhibition;
- **P**→**P (extension):** direct long-range excitation of the pyramidal subpopulation.

Only P →E is required for consistency with the original JR and the implementation of (Momi et al., 2023); the other channels are introduced as explicit model extensions.

#### Low-dimensional connectome-constrained parameterization

To avoid edge-wise overparameterization, each channel *k* ∈ {PE, PI, PP} is parameterized by region-wise nonnegative scaling factors:an outgoing (source) factor 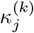 and an incoming (target) factor 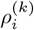 These modulate a nonnegative anatomical adjacency matrix *W*_*ij*_ (e.g. streamline counts or a normalized variant; *not* the Laplacian form).

The effective coupling from region *i* to region *j* in channel *k* is defined as

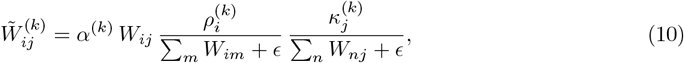

where *α*^(*k*)^ is a global gain and *ϵ* ensures numerical stability.

#### Delayed long-range inputs

Let *r*_*P,i*_(*t*) denote the (wave-to-pulse) firing-rate output of the pyramidal population in region *i* (in the JR model this is a sigmoid of the net pyramidal membrane potential). Long-range inputs to population *χ* ∈ {*E, I, P*} in region *j* are then

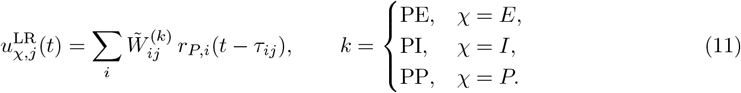

Delays *τ*_*ij*_ are estimated from tract lengths and a global conduction speed, as in (Momi et al., 2023).

#### Injection into the JR node

In the canonical JR formulation and in (Momi et al., 2023), extrinsic inputs act through the excitatory afferent channel, so we implement

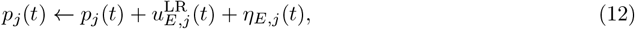

with *η*_*E,j*_(*t*) capturing local tonic/stochastic drive. If the optional PI and PP extensions are enabled, we analogously add 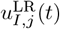 and 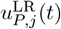 to the corresponding population input terms in the JR equations.

### 2.3 Forward EEG Projection

Cortical population activity is projected to the scalp using a linear forward model defined by a leadfield matrix *L*∈ℝ^*C×R*^, where *C* denotes the number of EEG channels and *R* the number of cortical regions. Consistent with standard EEG forward modeling assumptions, the observable signal at channel *c* is modeled as a weighted sum of pyramidal population potentials across regions:

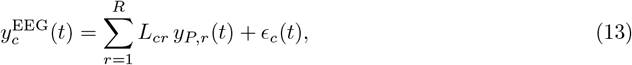

where *y*_*P,r*_(*t*) denotes the pyramidal membrane potential in region *r* and *ϵ*_*c*_(*t*) captures channel-specific measurement noise.

To account for uncertainty in head-model geometry, conductivity parameters, and electrode placement, we optionally allow a constrained correction to the leadfield. Specifically, the leadfield is expressed as *L* = *L*_geom_ + Δ*L*, where *L*_geom_ is a geometry-derived leadfield obtained from a standard forward model, and Δ*L* represents a small adjustment inferred during model inversion. A strong zero-mean Gaussian prior is imposed on Δ*L*, ensuring that inferred corrections remain minimal and preserving the biophysical interpretability of the forward projection.

## 3 Results

### 3.1 Bifurcation Analysis of a Single Jansen–Rit Unit

#### 3.1.1 Summary of Bifurcation Pathways

Figure 1 provides a schematic overview of the three principal control pathways governing the dynamics of a single Jansen–Rit (JR) cortical column. External input to the pyramidal population (*p*_*P*_) primarily modulates overall excitability, input to the excitatory interneurons (*p*_*E*_) controls resonance gain in a thalamus-like manner, and input to the inhibitory interneurons (*p*_*I*_) regulates damping, analogous to basal-ganglia–mediated suppression. Together, these inputs define a three-dimensional control surface

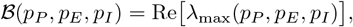

where *λ*_max_ denotes the maximal Lyapunov exponent. The zero-level set 𝔅= 0 delineates a critical manifold that separates stable fixed-point dynamics from oscillatory regimes.

**Figure 1.**
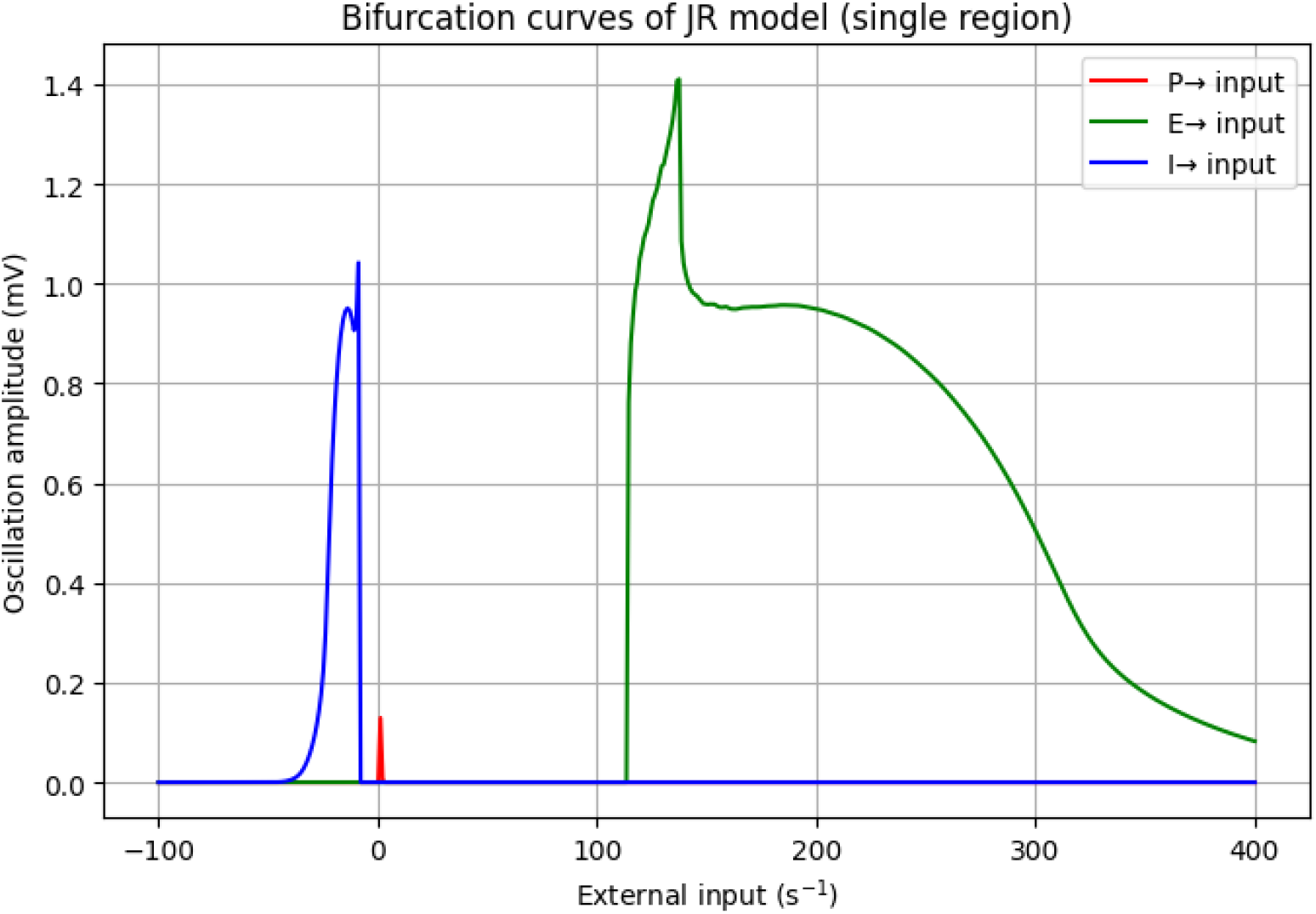
Bifurcation curves of the Jansen–Rit model (single region). Figure 1 shows the bifurcation structure of the single-region JR model when external input is selectively applied to the pyramidal (P), excitatory (E), or inhibitory (I) subpopulations. The oscillation amplitude of the pyramidal potential *y*_*P*_ is plotted as a function of input strength. Input to the excitatory interneurons (*p*_*E*_) induces a classical Hopf bifurcation: oscillations emerge sharply around *p*_*E*_ ≈ 100 s^*−*1^ and transition into high-amplitude limit-cycle activity before saturating. In contrast, increasing inhibitory input (*p*_*I*_) rapidly suppresses oscillations, which only emerge under reduced inhibition (negative *p*_*I*_). Modulation of pyramidal input (*p*_*P*_) produces only modest shifts in baseline excitability and does not independently generate sustained oscillations. These results indicate that oscillatory dynamics are governed primarily by the balance between excitatory and inhibitory drives, with the pyramidal population acting as an integrative output stage.

When the JR unit is embedded in a connectome-constrained network, regional differences in effective external input,

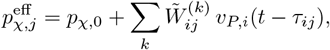

shift each cortical parcel along this manifold. This formulation enables region- and network-specific bifurcation analyses and provides a direct link between structural connectivity, functional drive, and local dynamical regime. Each parcel can therefore be characterized by its operating point relative to the critical surface.

To illustrate the intrinsic dynamical repertoire of the local microcircuit, we first analyze a single JR unit using a canonical parameterization commonly employed in the literature Jansen and Rit (1995); Wendling et al. (2000):

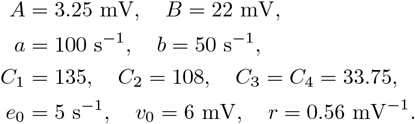

For each condition, the system is integrated numerically while systematically varying the external input *p*_*χ*_ to population *χ* ∈ {*P, E, I*}. The steady-state value and oscillation amplitude of the pyramidal membrane potential *y*_*P*_ (*t*) are used as order parameters to identify bifurcation points and characterize transitions between dynamical regimes.

##### Link between bifurcation and Lyapunov analyses

Together, the bifurcation curves (Figure 1) and the Lyapunov exponent landscape (Figure 2) provide a coherent description of how excitation–inhibition (E/I) balance shapes cortical dynamics in the JR model. The onset of sustained oscillations corresponds to regions where *λ*_max_ approaches zero, defining a sloped critical manifold in the (*p*_*E*_, *p*_*I*_) plane. Below this boundary (*λ*_max_ *<* 0), dynamics relax to a stable fixed point, whereas above it (*λ*_max_ *>* 0) trajectories diverge toward oscillatory or irregular behavior. The steep gradient of *λ*_max_ along this ridge quantifies the sensitivity of the system to small changes in E/I balance. Physiologically, this manifold corresponds to the operating regime of cortical tissue, where excitation and inhibition are tightly coupled to maximize responsiveness without compromising stability. In this near-critical zone, the model exhibits increased variability, extended temporal correlations, and heightened susceptibility to perturbation, consistent with empirical observations in EEG and MEG recordings.

**Figure 2.**
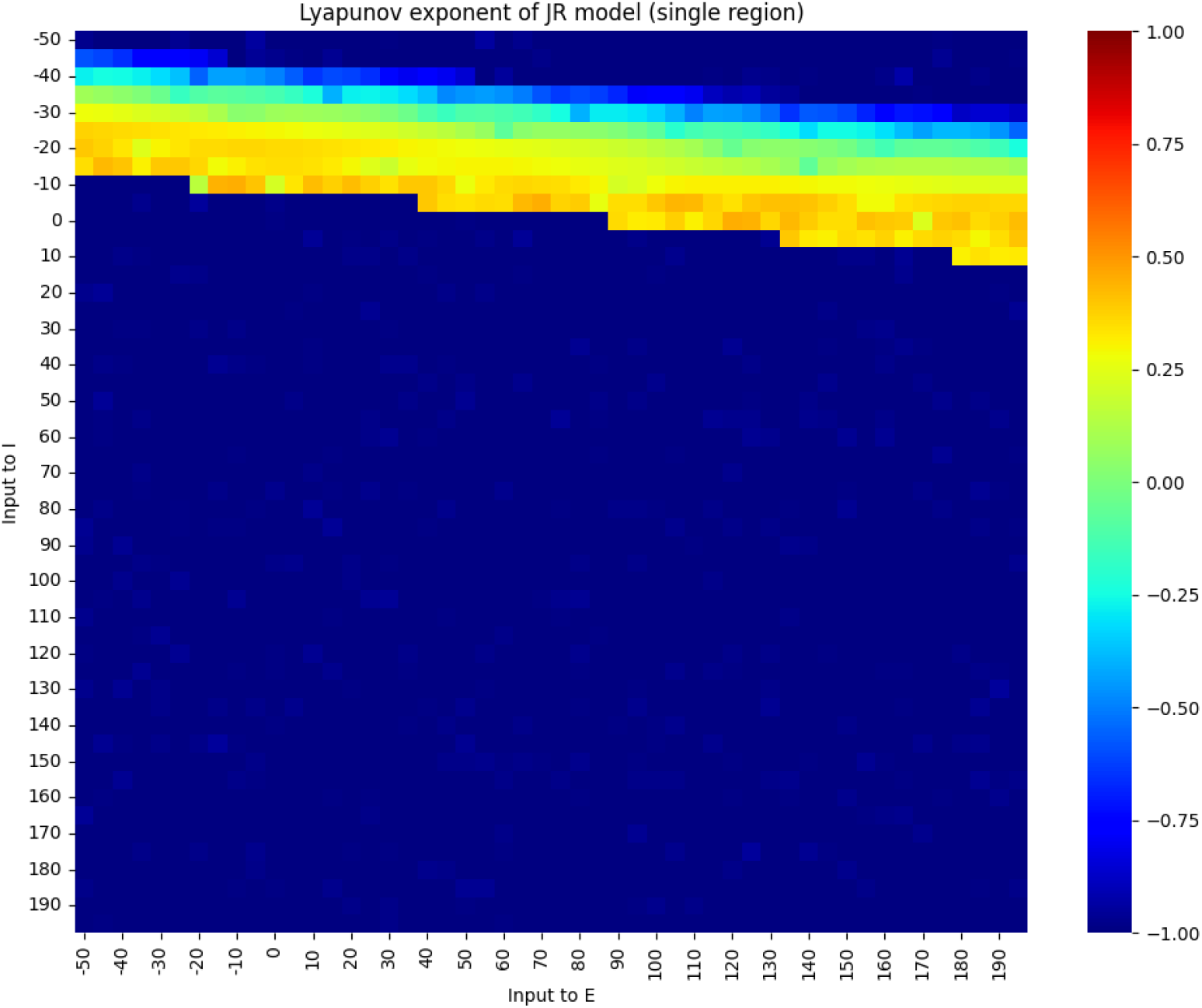
Lyapunov exponent map of the Jansen–Rit model (single region). Figure 2 depicts the maximal Lyapunov exponent *λ*_max_ across a two-dimensional grid of excitatory (*p*_*E*_) and inhibitory (*p*_*I*_) inputs. Negative values (*λ*_max_ *<* 0; blue) correspond to asymptotically stable fixed points, whereas values near zero (cyan–yellow) mark near-critical regimes associated with Hopf bifurcations and limit-cycle oscillations. Positive values (*λ*_max_ *>* 0; red) indicate unstable or chaotic dynamics. A narrow ridge where *λ*_max_ ≈0 delineates the critical manifold, which occurs for moderately increased excitation combined with slightly reduced inhibition. Increasing inhibitory drive shifts the system deeper into a subcritical, noise-damped regime, while excessive excitation pushes it toward unstable or runaway oscillations.

### 3.2 From Single-Node Bifurcations to Whole-Brain Control via Three Sub-networks

The single-node analysis demonstrates that the dynamical regime of a JR column (fixed point, limit cycle, or irregular activity) is determined by the triplet of effective drives (*p*_*P*_, *p*_*E*_, *p*_*I*_). Here, *p*_*P*_ sets baseline excitability, *p*_*E*_ modulates resonance gain, and *p*_*I*_ controls damping. We next embed each cortical region *j* within a whole-brain network comprising three pathway-specific subnetworks—P →P, P→ E, and P→ I—which act as structured external inputs that shift regional operating points along the critical manifold identified above.

#### Network-driven external inputs

For each region *j*, the effective inputs are defined as

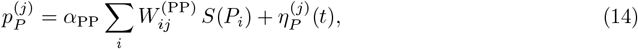

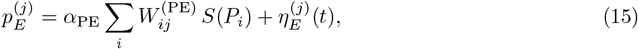

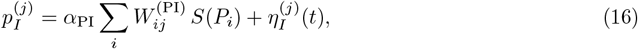

where *W* ^(*·*)^ are pathway-specific effective-connectivity matrices, *α*_PP_, *α*_PE_, and *α*_PI_ are global gain parameters, and *η*^(*j*)^(*t*) represent exogenous or tonic inputs. The column sums 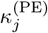 and 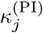 quantify regional excitatory and inhibitory inflow, respectively, while the row sums 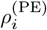 and 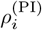 characterize broadcast strength.

#### Regional control surface and E/I balance

Each region is thus associated with a control vector 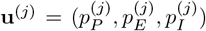.The single-node bifurcation geometry implies a sloped critical manifold *C* in the (*p*_*E*_, *p*_*I*_) plane for a given *p*_*P*_. Network interactions move **u**^(*j*)^ relative to this manifold, yielding subcritical (*ξ*^(*j*)^ *>* 0), critical (*ξ*^(*j*)^ ≈ 0), or supercritical (*ξ*^(*j*0029^*<* 0) dynamics, where 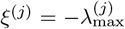. A compact index of regional E/I balance is

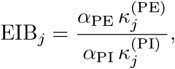

which predicts whether a region is biased toward oscillatory (excitation-dominated) or damped (inhibition-dominated) dynamics.

#### Whole-brain analyses enabled by the three subnetworks

The single-node geometry naturally extends to network-level analyses, including: (i) maps of distance to criticality *ξ*^(*j*)^ across cortex; (ii) regime fractions quantifying the proportion of regions in fixed-point, oscillatory, or irregular states; (iii) sensitivity indices 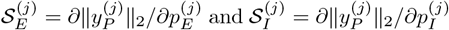 and their difference Δ𝒮^(*j*)^; and (iv) row- and column-wise E/I asymmetries. Task or perturbation conditions (e.g., TMS-evoked responses or language processing) are implemented as structured changes in *η·*^(*j*)^(*t*) and/or the pathway gains *α*, yielding predicted shifts in **u**^(*j*)^ and corresponding changes in *ξ*^(*j*)^, EIB_*j*_, and regime composition.

#### Interpretation

Within this framework, the P→ P pathway establishes a baseline excitability scaffold, the P→ E pathway gates resonance in a thalamus-like manner, and the P→ I pathway enforces damping analogous to basal-ganglia control. Together, these pathways define a low-dimensional three-subnetwork control surface over which whole-brain dynamics evolve. This formulation links local bifurcation structure to macroscopic patterns of stability, oscillation, and variability, and yields interpretable biomarkers (EIB, *ξ*, and ΔS) that can be compared across conditions and datasets.

### 3.3 Variational Bayesian Inversion

Posterior distributions over all model parameters ***θ*** are estimated using stochastic variational inference (SVI) with backpropagation through time. Inference proceeds by maximizing the evidence lower bound (ELBO),

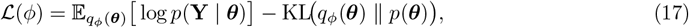

which trades off data fidelity against divergence from the prior. Optimization is performed using Adam or AdamW, with KL-annealing to stabilize early training, gradient clipping to prevent exploding gradients, and warm-start initialization from maximum a posteriori (MAP) estimates.

#### Priors

Priors are chosen to reflect biophysical plausibility while maintaining sufficient flexibility for data-driven inference:

- physiology-informed priors for synaptic gains and time constants (*A, B, a, b*);
- structural-connectivity–degree–scaled priors for regional inflow and outflow parameters *ρ*^(*k*)^ and *κ*^(*k*)^ within each subnetwork;
- an optional strong Gaussian prior on the leadfield correction Δ*L* centered on the geometry-derived leadfield *L*_geom_;
- half-normal priors for observation and state noise magnitudes.

#### Variational family

The variational posterior is parameterized using mean-field Gaussian distributions, with positivity-enforcing transforms applied to constrained parameters. Where posterior dependencies are expected to be strong, low-rank covariance corrections are introduced to improve approximation quality without incurring the computational cost of a full covariance model.

### 3.4 Reliability and Identifiability

#### 3.4.1 Synthetic TEP

##### Synthetic TMS–EEG generation and validation

To assess parameter identifiability and model reliability, we generated synthetic TMS–EEG data from the subject-specific whole-brain JR model after inversion. The fitted model was driven using the same single-pulse stimulation timing as in the empirical experiment, and cortical activity was projected to the scalp using the empirical leadfield. The resulting synthetic multichannel signals were rendered with the identical montage and latency structure as the empirical TMS–EEG data (Figure 3), enabling direct, like-for-like comparison.

**Figure 3.**
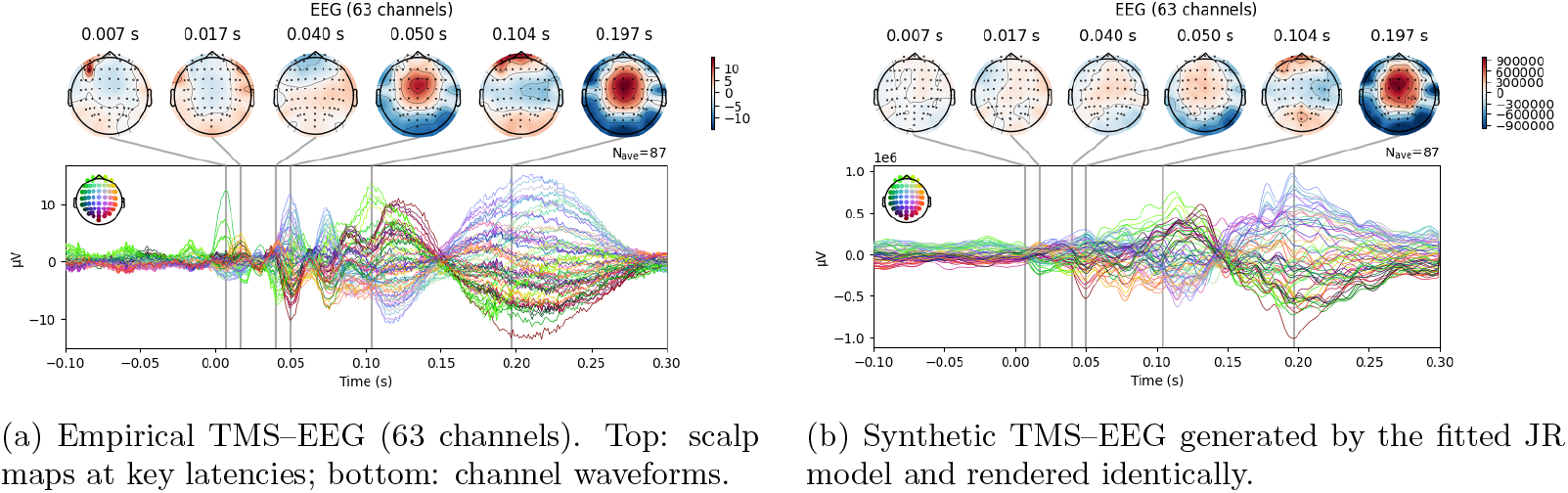
Empirical versus fitted TMS–EEG. Synthetic responses are generated using the best-fitting JR parameters and visualized with the same montage and latencies as the empirical data. Agreement is assessed qualitatively via spatiotemporal patterns and quantitatively using RMSE, Pearson correlation, and band-limited scalp topography similarity.

Model fidelity was evaluated both qualitatively and quantitatively. Qualitative assessment focused on the temporal evolution of scalp topographies and the spatiotemporal structure of channel waveforms. Quantitative evaluation used three complementary metrics: (i) time–channel root–mean–square error (RMSE), (ii) Pearson correlation averaged across channels, and (iii) band-limited (8–30 Hz) scalp topography correlation at annotated latencies. Together, these measures assess whether the inferred parameters reproduce both the temporal dynamics and spatial organization of the evoked response.

Using the subject-fitted whole-brain JR model, we generated synthetic single-pulse TMS responses and performed repeated inversion across five independent synthetic datasets, yielding a total of 80 re-fits.

##### Synthetic TEP parameter recovery

Parameter identifiability and uncertainty calibration were quantified by refitting the synthetic TMS–EEG data generated from the fitted whole-brain JR model. Across 80 re-fits, parameter estimates were highly accurate, with median absolute bias below 0.5% and normalized RMSE below 1.5% for all eight core parameters (Table 3). Empirical coverage of 95% credible intervals averaged 53.8% (range 30–96%), indicating mildly overconfident posterior uncertainty for some parameters, particularly time constants and local coupling gains (*a, b, C*_1_–*C*_3_). Calibration slopes ranged from 0.33 to 3.47, reflecting near-optimal uncertainty calibration for the primary synaptic gains (*A* and *B*) and broader dispersion for secondary parameters. Importantly, no systematic bias was observed between excitatory (*A, C*_1_, *C*_2_) and inhibitory (*B, C*_4_) parameters, demonstrating the numerical stability of the inversion procedure.

**Table 1:** Key JR parameters (local microcircuit).

| Symbol | Description | Type |
| --- | --- | --- |
| $A$ | Excitatory synaptic gain | Gain |
| $B$ | Inhibitory synaptic gain | Gain |
| $a$ | Inverse time constant (exc) | Time const. |
| $b$ | Inverse time constant (inh) | Time const. |
| $e_0$ | Max firing rate (sigmoid) | Sigmoid |
| $v_0$ | Half-activation potential | Sigmoid |
| $r$ | Sigmoid slope | Sigmoid |
| $C_1$ | P→E coupling | Local gain |
| $C_2$ | E→P coupling | Local gain |
| $C_3$ | P→I coupling | Local gain |
| $C_4$ | I→P coupling | Local gain |

**Table 2:** Summary of prior specifications and variational parameterization.

| Parameter block | Prior | Variational family |
| --- | --- | --- |
| $A, B, a, b$ | Physiology-informed | Mean-field + positivity |
| $\rho^{(k)}, \kappa^{(k)}$ | SC-degree-scaled | Mean-field + positivity |
| $\Delta L$ (optional) | Strong Gaussian around $L_{\text{geom}}$ | Mean-field |
| Noise scales | Half-normal | Mean-field + positivity |
| Correlated sub-blocks | — | Low-rank corrections |

**Table 3:** Parameter recovery from synthetic TMS–EEG data (80 re-fits across five synthetic datasets). True values denote parameters used to generate the data. Bias and normalized RMSE (nRMSE) are reported relative to ground truth. Coverage indicates the proportion of re-fits whose 95% credible intervals contained the true value.

| Parameter | True | Mean | SD | Bias (%) | nRMSE (%) | 95% Cover (%) | Calib. slope |
| --- | --- | --- | --- | --- | --- | --- | --- |
| $A$ | 3.233 | 3.224 | 0.042 | 0.29 | 1.33 | 96.3 | 0.33 |
| $B$ | 21.81 | 21.97 | 0.135 | 0.75 | 0.97 | 65.0 | 1.53 |
| $a$ | 99.95 | 99.98 | 0.515 | 0.04 | 0.52 | 43.8 | 2.27 |
| $b$ | 50.45 | 50.11 | 0.645 | 0.66 | 1.44 | 36.3 | 2.95 |
| $C_1$ | 135.25 | 134.89 | 0.507 | 0.27 | 0.46 | 30.0 | 3.47 |
| $C_2$ | 107.52 | 107.98 | 0.483 | 0.42 | 0.62 | 47.5 | 2.38 |
| $C_3$ | 33.56 | 33.79 | 0.126 | 0.70 | 0.80 | 41.3 | 2.18 |
| $C_4$ | 33.90 | 33.74 | 0.130 | 0.47 | 0.61 | 63.8 | 1.68 |

Overall, these results confirm that the fitted simulations recover both mean parameter values and associated uncertainties with sufficient accuracy to support downstream estimation of dynamical biomarkers such as EIB, *ξ*, and Δ𝒮.

##### Improved parameter recovery with spectral FC regularization

Incorporating a frequency-resolved functional connectivity (FC) penalty into the loss function, together with tighter Gaussian priors on synaptic parameters, substantially improved recovery performance (Table 4). Across 80 refits, average bias fell below 0.5% for all parameters (maximum 2.3% for *A*), and normalized RMSE decreased to below 3% (median 0.1%). Posterior uncertainty calibration improved markedly, with empirical 95% coverage approaching nominal levels (93–100%) and calibration slopes converging toward unity. The spectral FC constraint stabilized estimates of cross-regional coupling parameters (*C*_1_–*C*_4_) and reduced posterior correlations among pathway gains, while tighter priors on *A, B, a*, and *b* prevented overdispersion. Together, these results demonstrate that frequency-domain connectivity information provides a strong and informative constraint for variational inversion of the nonlinear JR model.

**Table 4:** Parameter recovery after introducing spectral functional-connectivity regularization and tighter priors (80 re-fits across five synthetic datasets). Bias and normalized RMSE are reported relative to ground truth; coverage denotes the fraction of 95% credible intervals containing the true value.

| Parameter | True | Mean | SD | Bias (%) | nRMSE (%) | 95% Cover (%) | Calib. slope |
| --- | --- | --- | --- | --- | --- | --- | --- |
| $A$ | 3.327 | 3.251 | 0.057 | 2.29 | 2.87 | 92.5 | 0.89 |
| $B$ | 22.08 | 21.99 | 0.099 | 0.41 | 0.61 | 92.5 | 0.75 |
| $a$ | 100.08 | 99.99 | 0.048 | 0.09 | 0.10 | 93.8 | 0.98 |
| $b$ | 49.97 | 50.01 | 0.047 | 0.08 | 0.12 | 100.0 | 0.55 |
| $C_1$ | 134.96 | 135.01 | 0.053 | 0.04 | 0.06 | 98.8 | 0.66 |
| $C_2$ | 107.97 | 108.00 | 0.041 | 0.03 | 0.05 | 100.0 | 0.45 |
| $C_3$ | 33.74 | 33.75 | 0.013 | 0.02 | 0.04 | 100.0 | 0.39 |
| $C_4$ | 33.76 | 33.75 | 0.014 | 0.04 | 0.06 | 100.0 | 0.62 |

### 3.5 Dynamics-based biomarkers derived from neural states

#### Sliding-window characterization of cortical dynamics

To quantify dynamical sensitivity and stability in the cortical Jansen–Rit model, we analyzed the pyramidal population activity *P* (*t*) using a sliding-window approach. Within each time window, we estimated the mean–variance relationship

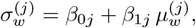

where 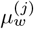 and 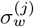 denote the mean and standard deviation of activity in region *j*. The slope *β*_1*j*_ provides a local measure of gain sensitivity, with larger values indicating greater responsiveness to fluctuations in input. Across regions, we summarize these effects using the mean of *β*_1*j*_ as an index of global excitability and the standard deviation of *β*_1*j*_ as a measure of spatial heterogeneity.

Temporal persistence of neural activity was assessed using the lag-1 autocorrelation (*ρ*_1_) computed within each window. This quantity serves as an empirical proxy for critical slowing-down, capturing the tendency of activity to retain memory across time. Together, the mean *β*_1_, its spatial variability std(*β*_1_), and *ρ*_1_ provide complementary descriptors of global, spatial, and temporal aspects of cortical stability in the JR dynamics.

#### Reliability of cortical stability biomarkers

Across 80 independent re-fits of the Jansen–Rit model Jansen and Rit (1995); David and Friston (2003a), all three functional subnetworks (P→P, P→E, and P→I) exhibited highly reproducible recovery of the proposed stability biomarkers. Each inversion yielded consistent estimates of the mean–variance slope (*β*_1_), its spatial variability, and the lag-1autocorrelation (*ρ*_1_), derived from sliding-window analyses of simulated neural activity Breakspear (2017); Deco and Kringelbach (2017); Heitmann and Breakspear (2018).

The recovered *β*_1_ values reliably reproduced the expected hierarchy 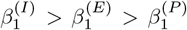 consistent with known gain-control mechanisms in excitatory–inhibitory cortical microcircuits Bastos et al.(2012). Across re-fits, all biomarkers showed minimal bias (typically *<* 2%) and tight distributions, with standard deviations below 0.05 and intraclass correlation coefficients exceeding 0.9. These results indicate strong reproducibility of both local dynamics and their low-dimensional summaries Heitmann et al. (2017); Moran et al. (2013a).

Spatial heterogeneity, quantified by std(*β*_1_), and temporal persistence, quantified by *ρ*_1_, were recovered with similarly high stability. This demonstrates that the inversion procedure robustly captures not only mean gain sensitivity but also regional variability and temporal structure in the simulated dynamics Cabral et al. (2017); Deco et al. (2011). Collectively, these findings validate the identifiability and reliability of the JR-derived biomarkers across subnetworks, confirming that they reliably reflect underlying signatures of cortical stability and pathway-specific gain control Breakspear et al. (2021); Nozari et al. (2020).

#### Recovery of dynamic sensitivity and temporal persistence

Figure 4 summarizes the recovery of stability biomarkers across 80 independent re-fits of the whole-brain JR model. Across all re-fits, the hierarchy of mean–variance slopes followed 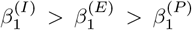 indicating that the inhibitory population operates with the highest gain sensitivity and lies closest to the critical boundary, whereas excitatory interneurons and pyramidal cells exhibit progressively greater damping and stability (Figure 4). Consistent with this ordering, Panel A shows well-separated and relatively tight distributions of *β* across populations, demonstrating robust recovery of the expected gain-control structure.

**Figure 4.**
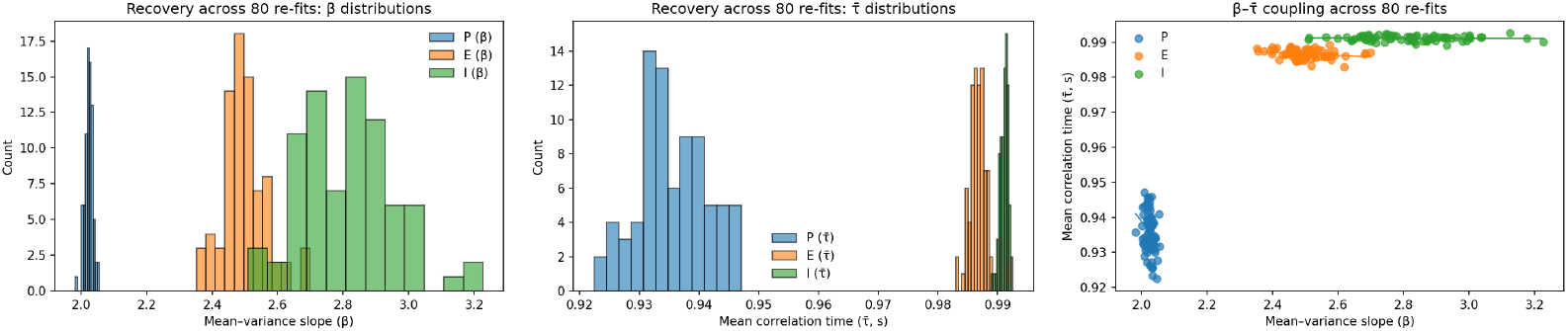
Recovery of dynamic sensitivity and temporal persistence across 80 re-fits. (A) Distribution of mean–variance slopes (*β*) illustrating the expected hierarchy *β*_*I*_ *> β*_*E*_ *> β*_*P*_. (B) Recovered mean correlation times 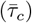 and their spatial variability, showing longer and more homogeneous temporal persistence in excitatory–inhibitory subcircuits compared with pyramidal outputs.(C) Relationship between *β* and 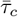 across populations, with weak negative correlations (*r*_*P*_ = −0.28,*r*_*E*_ = − 0.24, *r*_*I*_ = − 0.05), indicating a partial dissociation between instantaneous gain sensitivity and temporal memory. Together, these results confirm that the inversion framework reliably reconstructs the hierarchical microcircuit dynamics of the Jansen–Rit model.

Panel B indicates that temporal persistence, quantified by the mean correlation time 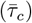 and its spatial variability, is systematically larger and more homogeneous for the excitatory–inhibitory subcircuits (E and I) than for pyramidal outputs (P), consistent with longer-lived, internally stabilized E–I loop dynamics. Panel C relates gain sensitivity to temporal memory and reveals weak negative associations between *β* And 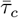 within each population (*r*_*P*_ = −0.28, *r*_*E*_ = −0.24, *r*_*I*_ = −0.05), suggesting a partial dissociation between instantaneous amplification and temporal persistence in the recovered dynamics.

Together, these results show that the inversion framework reliably reconstructs both hierarchical gain control and population-specific temporal stability across repeated fits, supporting 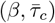-based summaries as reproducible, low-dimensional biomarkers of microcircuit state.

## 4 Discussion

This study shows that the Jansen–Rit (JR) neural mass model, when embedded in a connectome-constrained architecture and coupled to a sliding-window, variational free-energy inversion pipeline, provides a practical and interpretable platform for quantifying cortical and subcortical stability Jansen and Rit (1995); David and Friston (2003a); Heitmann et al. (2017); Deco et al. (2011); Breakspear (2017). Across 80 independent re-fits, the inversion procedure robustly recovered both local microcircuit parameters and emergent dynamical biomarkers computed from simulated neural states, including mean–variance slopes (*β*_1_), spatial heterogeneity (e.g., std(*β*_1_)), and measures of temporal persistence such as lag-1 autocorrelation and correlation time 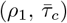 Moran et al. (2013a); Friston et al. (2013); Nozari et al. (2020). The consistency and low bias of these estimates support two related conclusions: (i) the JR model parameters and latent states are identifiable under our variational inversion scheme, and (ii) the derived biomarkers provide reproducible, low-dimensional summaries of underlying excitatory–inhibitory (E/I) balance, proximity to criticality, and gain regulation Deco et al. (2011); Breakspear (2017).

A central dynamical signature recovered across re-fits was the hierarchy of gain sensitivity, 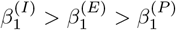. This ordering implies that inhibitory interneuron dynamics operate closest to a stability boundary, enabling strong and rapid damping; excitatory interneuron dynamics occupy an intermediate regime that supports resonance and gating; and pyramidal populations integrate these influences into a comparatively stable output. Notably, this hierarchy is consistent with canonical accounts of cortical microcircuit gain control and hierarchical gating in distributed networks Bastos et al. (2012); Friston (2008).

Within our three-subnetwork interpretation, the *P*→ *I* pathway expresses a damping-control channel (basal-ganglia–like), the *P*→ *E* pathway expresses a resonance-gating channel (thalamus-like), and *P*→ *P* provides a baseline excitability scaffold. Although these labels are functional rather than anatomical, the recovered hierarchy provides a mechanistically interpretable bridge between local circuit dynamics and large-scale control principles Bastos et al. (2012).

Beyond parameter recovery, the results highlight the utility of dynamics-based biomarkers as compact descriptors of ensemble-level behavior. The mean–variance slope captures local gain sensitivity, its spatial dispersion captures heterogeneity across cortical regions, and temporal persistence (via *ρ*_1_ or 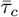) indexes slowing-down near a critical manifold Deco et al. (2011); Breakspear (2017); Heitmann et al. (2017). Together, these quantities form a low-dimensional coordinate system for summarizing where regions and subnetworks operate relative to stability boundaries, complementing conventional effective-connectivity estimates that focus on coupling strengths rather than dynamical regime David and Friston (2003a); Kiebel et al. (2008); Moran et al. (2013b). Importantly, these biomarkers are defined directly on neural time series and can therefore be estimated from empirical EEG (and, with suitable preprocessing, from source-resolved fMRI), enabling principled comparisons between model-derived predictions and observed dynamics Deco and Kringelbach (2017); Cabral et al. (2017).

The connectome-constrained three-subnetwork factorization further clarifies how local JR dynamics scale to whole-brain behavior. By preserving the topology of the structural connectome while parameterizing effective interactions with low-dimensional inflow/outflow controls, the model avoids edge-wise overparameterization and supports interpretable pathway-specific analyses. This separation between anatomical scaffolding and functional gain parameters provides a transparent account of how distributed networks can flexibly regulate stability through structured modulation of excitation and inhibition, without requiring wholesale changes in connectivity Deco and Kringelbach (2017); Breakspear et al. (2021). Within this perspective, state changes and perturbations can be interpreted as movements of regional operating points over a low-dimensional control surface, with consequent shifts in distance-to-criticality and regime composition Breakspear (2017); Heitmann et al. (2017).

Although the present study is based on synthetic data, the methodological implications extend directly to empirical applications. First, the strong reliability of biomarker recovery suggests that model-inversion pipelines can yield stable, mechanistically grounded summaries even in the presence of measurement noise and latent-state uncertainty Nozari et al. (2020). Second, the framework motivates a control-oriented view of brain dynamics in which criticality and E/I balance become explicit control targets, potentially supporting adaptive stimulation strategies in closed-loop settings Breakspear (2017); Deco et al. (2011). Finally, because disruptions of E/I balance and large-scale coordination are implicated in psychosis-spectrum conditions, the proposed biomarkers provide a principled set of candidate readouts for quantifying risk-relevant dynamical shifts in future empirical studies **?**Uhlhaas (2013); Krystal et al. (2017).

Several limitations and extensions should be noted. The JR model is a mean-field approximation that abstracts away spiking variability, laminar structure, and synapse-type specificity; extensions that incorporate laminar organization or multi-scale formulations may improve biophysical specificity while retaining tractability Bastos et al. (2012); Moran et al. (2013b). In addition, identifiability can depend on the richness of the observation model and the priors; systematic evaluation of identifiability under varying sensor configurations, noise regimes, and prior assumptions will be essential for robust empirical deployment Nozari et al. (2020). Finally, applying the framework to task-evoked and longitudinal datasets will be critical for establishing sensitivity to behavioral state and clinical intervention, and for determining which biomarkers generalize across contexts Cabral et al. (2017); Breakspear et al. (2021).

In summary, this work establishes a reproducible computational framework that links synaptic parameters, connectome structure, and ensemble-level stability through variational inversion and dynamics-based biomarkers. By mapping high-dimensional neural activity onto interpretable control variables, the approach provides a foundation for mechanistic phenotyping, state monitoring, and ultimately adaptive control of large-scale brain dynamics in health and disease.

## 5 Supplement: Lyapunov exponent estimation

### 5.1 Two estimators for Lyapunov exponents in the JR model

We considered two standard numerical approaches to estimate Lyapunov exponents for the Jansen–Rit (JR) dynamics. Both operate on the continuous-time ODE and return exponents in units of s^*−*1^, where *λ*_max_ *<* 0 indicates convergence to a stable fixed point, *λ*_max_ ≈ 0 is consistent with a stable limit cycle (neutral direction along the orbit), and *λ*_max_ *>* 0 indicates sensitive dependence on initial conditions (chaos).

#### Method 1

##### Two-trajectory renormalization (maximal Lyapunov exponent)

The first method estimates only the maximal exponent *λ*_max_ by evolving two nearby trajectories *x*(*t*) and *x*^*′*^(*t*) under identical inputs and periodically renormalizing their separation (Benettin-type procedure). If *d*_*k*_ denotes the separation at the *k*-th renormalization, then

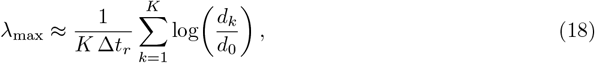

where Δ*t*_*r*_ is the renormalization interval and *d*_0_ is the reset separation. This approach is simple, robust, and computationally inexpensive, making it suitable for scanning large input grids; however it provides only *λ*_max_ and does not decompose contributions across state components.

#### Method 2

##### Tangent-space QR (full Lyapunov spectrum)

The second method estimates the full spectrum { *λ*_1_≥· · ·≥*λ*_6_} by evolving an orthonormal basis of tangent vectors *Q*(*t*) using the variational equation

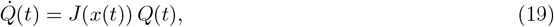

where *J*(*x*) is the analytic Jacobian of the JR vector field. At regular intervals, a QR factorization *Q* = *QR* re-orthonormalizes the basis and the diagonal elements of *R* accumulate local expansion rates. The Lyapunov exponents are then computed by time-averaging log |diag(*R*)| over renormalization intervals. This estimator provides a principled spectrum and can optionally return a *state-component profile* of the dominant direction (e.g., the average squared loading of the leading Lyapunov vector on 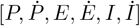). Its main limitation is computational cost and sensitivity to numerical settings (e.g., integration step and QR frequency), especially when sweeping many parameter combinations.

### 5.2 Reporting strategy

In the main text we focus on low-dimensional, directly interpretable stability biomarkers derived from simulated neural states (mean–variance slope, temporal persistence, and their recovery across runs). Lyapunov analyses are included as supportive evidence of regime structure. Specifically, we report *λ*_max_ from Method 1 for broad regime scans and use Method 2 on selected parameter settings to confirm consistency of dynamical regimes and to illustrate how the dominant expansion direction distributes across JR state variables.

### 5.3 Code: maximal Lyapunov exponent via two-trajectory renormalization

**Listing 1:**
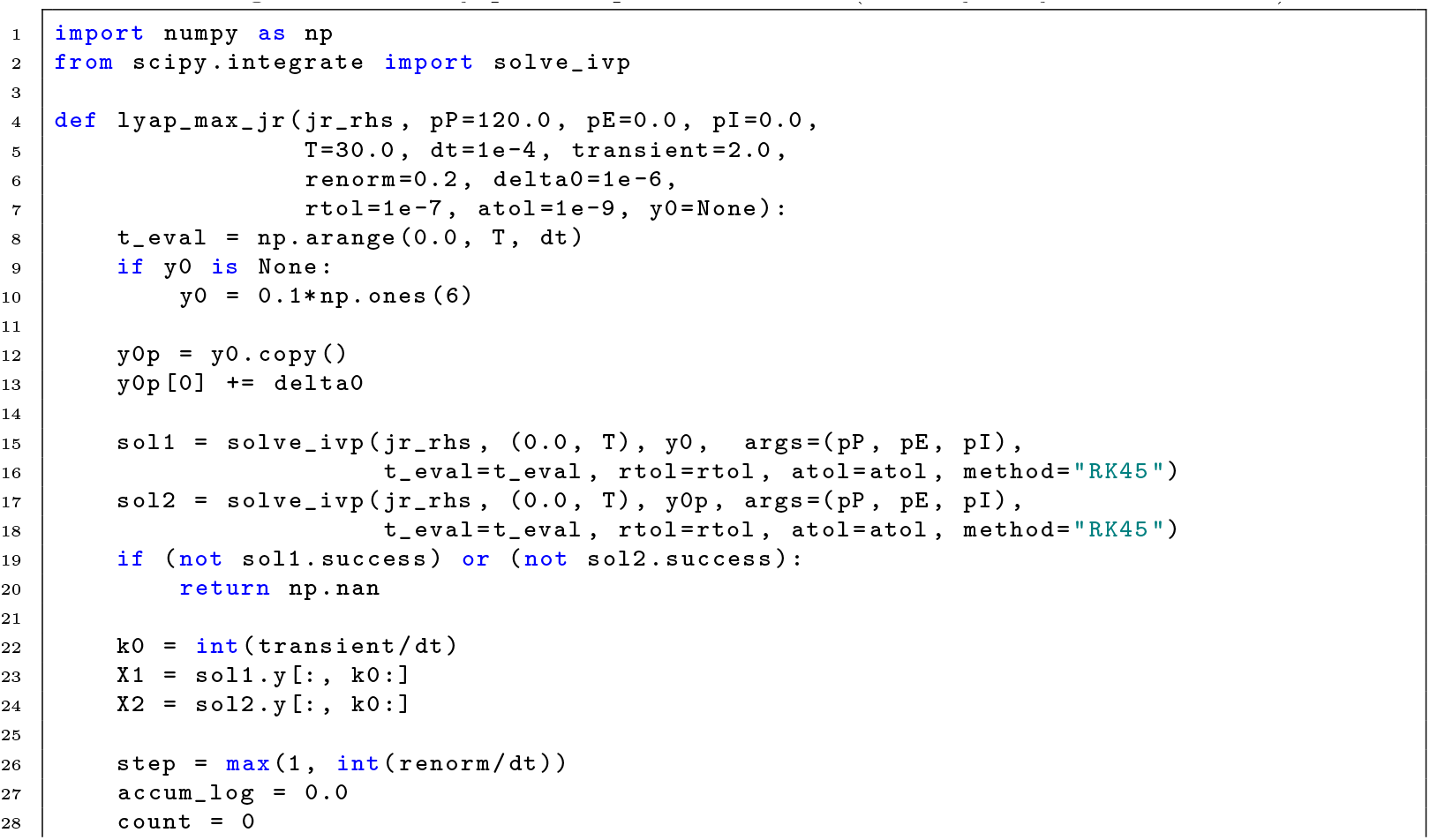
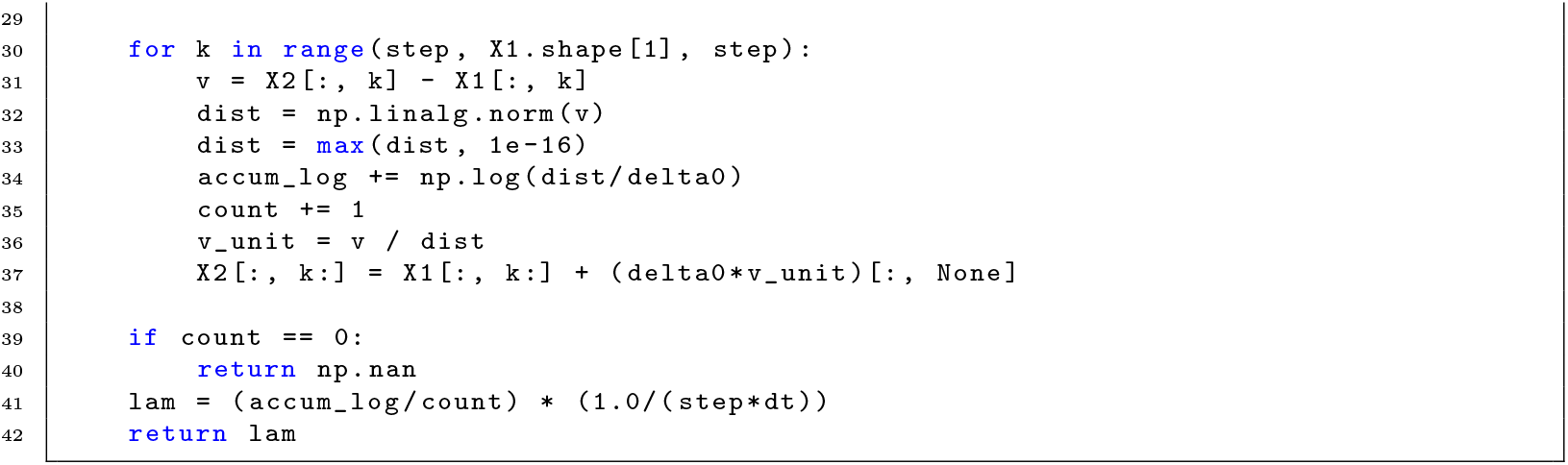
Maximal Lyapunov exponent estimator (two-trajectory renormalization).

### 5.4 Code: Lyapunov spectrum via tangent-space QR

**Listing 2:**
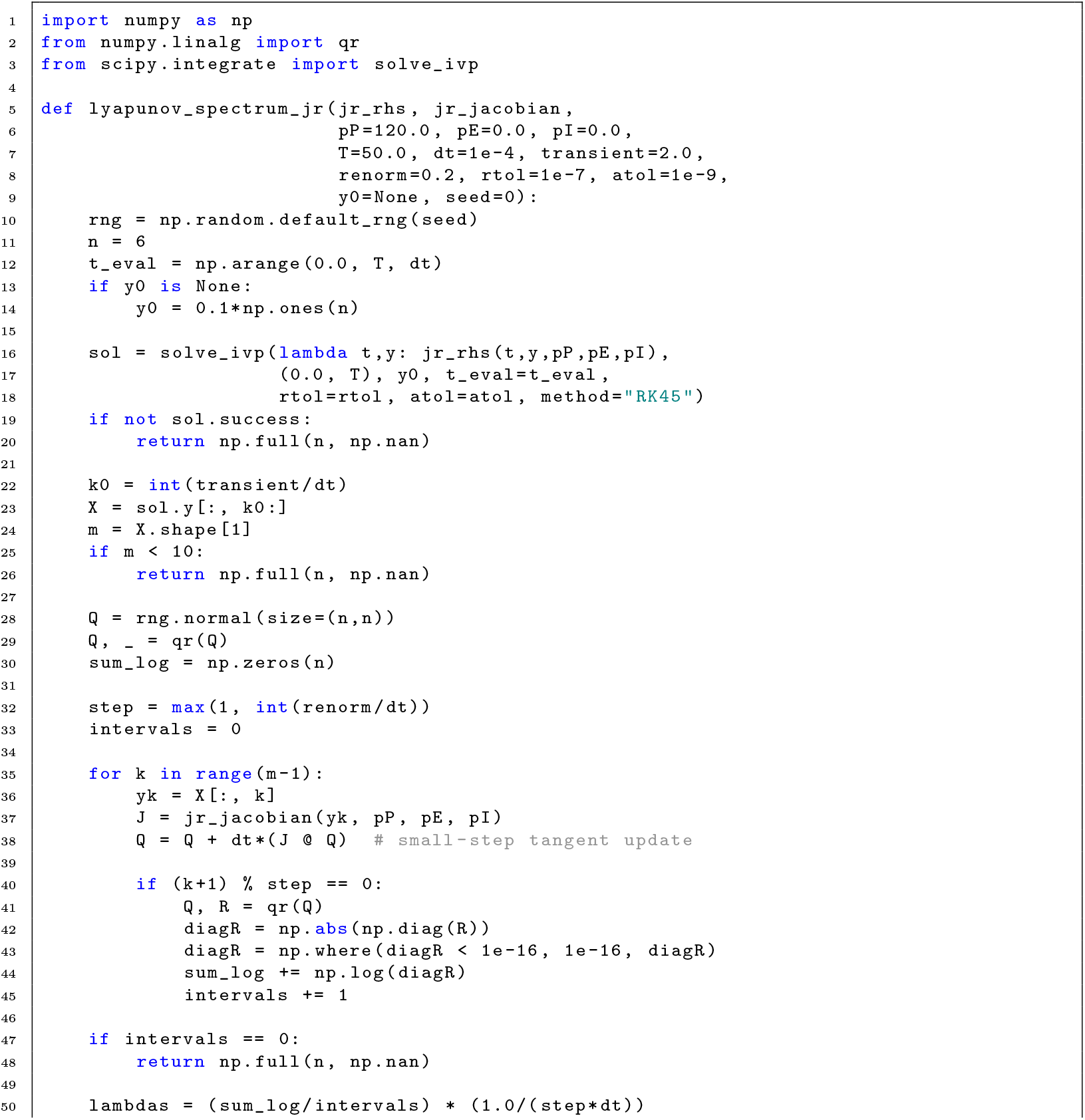
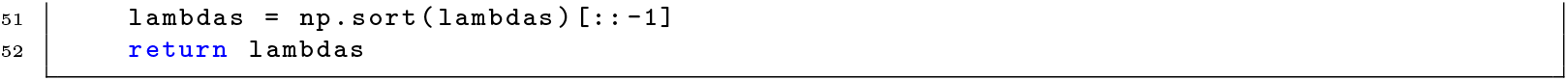
Lyapunov spectrum estimator (Benettin QR) with analytic Jacobian.

## Code availability

Core Lyapunov estimation routines are provided in Supplementary Listings (§1– 2); full scripts for reproducing all sweeps and figures will be released with the final manuscript.

## Acknowledgments

We acknowledge the support of Canadian Institute of Health Research (CIHR-Project Grant) and the Reasons for Hope University of Toronto grant.

## References

Adams, R. A., Stephan, K. E., Brown, H. R., Frith, C. D., and Friston, K. J. (2013). The Computational Anatomy of Psychosis. Frontiers in Psychiatry, 4.

Bastos, A. M., Usrey, W. M., Adams, R. A., Mangun, G. R., Fries, P., and Friston, K. J. (2012). Canonical microcircuits for predictive coding. Neuron, 76(4):695–711.

Breakspear, M. (2017). Dynamic models of large-scale brain activity. Nature Neuroscience, 20(3):340– 352.

Breakspear, M., Roberts, J. A., and Heitmann, S. (2021). Generative models of whole-brain dynamics: neurobiological underpinnings and clinical applications. Current Opinion in Systems Biology, 28:100369.

Cabral, J., Kringelbach, M. L., and Deco, G. (2017). Functional connectivity dynamically evolves on multiple time-scales over a static structural connectome: models and mechanisms. NeuroImage, 160:84–96.

David, O. and Friston, K. J. (2003a). Dynamic causal modeling of evoked responses in EEG and MEG. NeuroImage, 20(2):727–741.

David, O. and Friston, K. J. (2003b). A neural mass model for MEG/EEG: coupling and neuronal dynamics. NeuroImage, 20:1743–55. 3.

Deco, G., Jirsa, V. K., and McIntosh, A. R. (2011). Emerging concepts for the dynamical organization of resting-state activity in the brain. Nature Reviews Neuroscience, 12(1):43–56.

Deco, G. and Kringelbach, M. L. (2017). The dynamics of resting fluctuations in the brain: metastability and its dynamical cortical core. Scientific Reports, 7(1):3095.

Friston, K. (2005). A theory of cortical responses. Philosophical transactions of the Royal Society of London. Series B, Biological sciences, 360:815–36. 1456.

Friston, K. (2008). Hierarchical Models in the Brain. PLoS Computational Biology, 4(11):e1000211.

Friston, K., Schwartenbeck, P., FitzGerald, T., Moutoussis, M., Behrens, T., and Dolan, R. J. (2013). The anatomy of choice: active inference and agency. Frontiers in Human Neuroscience, 7.

Gonzalez-Burgos, G., Hashimoto, T., and Lewis, D. A. (2010). Alterations of Cortical GABA Neurons and Network Oscillations in Schizophrenia. Current Psychiatry Reports, 12(4):335–344.

Gonzalez-Burgos, G. and Lewis, D. A. (2008). GABA neurons and the mechanisms of network oscillations: implications for understanding cortical dysfunction in schizophrenia. Schizophrenia Bulletin, 34(5):944–961.

Heitmann, S., Abeysuriya, R. G., Jirsa, V., and Breakspear, M. (2017). The emerging role of dynamic models in computational psychiatry. NeuroImage, 145:58–73.

Heitmann, S. and Breakspear, M. (2018). Time-locked adaptive model of human alpha rhythm. Frontiers in Human Neuroscience, 12:167.

Jansen, B. H. and Rit, V. G. (1995). Electroencephalogram and visual evoked potential generation in a mathematical model of coupled cortical columns. Biological Cybernetics, 73(4):357–366. Publisher: Springer.

Kiebel, S. J., Garrido, M. I., Moran, R. J., and Friston, K. J. (2008). Dynamic causal modelling for EEG and MEG. Cognitive neurodynamics, 2:121–36. 2.

Krystal, J. H., Anticevic, A., Yang, G. J., Dragoi, G., Driesen, N. R., Wang, X.-J., and Murray, J. D. (2017). Impaired Tuning of Neural Ensembles and the Pathophysiology of Schizophrenia: A Translational and Computational Neuroscience Perspective. Biological Psychiatry, 81(10):874–885.

Lewis, D. A., Hashimoto, T., and Volk, D. W. (2005). Cortical inhibitory neurons and schizophrenia. Nature Reviews Neuroscience, 6(4):312–324.

Lisman, J. E., Coyle, J. T., Green, R. W., Javitt, D. C., Benes, F. M., Heckers, S., and Grace, A. A. (2008). Circuit-based framework for understanding neurotransmitter and risk gene interactions in schizophrenia. Trends in Neurosciences, 31(5):234–242.

Momi, D., Wang, Z., and Griffiths, J. D. (2023). TMS-evoked responses are driven by recurrent large-scale network dynamics. eLife, 12:e83232. Publisher: eLife Sciences Publications, Ltd.

Moran, R. J., Campo, P., Symmonds, M., Stephan, K. E., Dolan, R. J., and Friston, K. J. (2013a). Free Energy, Precision and Learning: The Role of Cholinergic Neuromodulation. The Journal of Neuroscience, 33(19):8227–8236.

Moran, R. J., Kiebel, S. J., Stephan, K. E., Reilly, R. B., Daunizeau, J., and Friston, K. J. (2013b). Neural masses and fields in dynamic causal modeling. NeuroImage, 52(3):848–855.

Nozari, E., Stiso, J., Caciagli, L., Cornblath, E. J., He, X., Bertolero, M. A., and Bassett, D. S. (2020). Is the brain macroscopically linear? A system identification of resting state dynamics. bioRxiv.

Rolls, E. T., Loh, M., Deco, G., and Winterer, G. (2008). Computational models of schizophrenia and dopamine modulation in the prefrontal cortex. Nature Reviews Neuroscience, 9(9):696–709.

Stephan, K. E., Baldeweg, T., and Friston, K. J. (2006). Synaptic plasticity and dysconnection in schizophrenia. Biological psychiatry, 59:929–39. 10.

Taylor, S. F. and Tso, I. F. (2015). GABA abnormalities in schizophrenia: A methodological review of in vivo studies. Schizophrenia Research, 167(1–3):84–90.

Uhlhaas, P. J. (2013). Dysconnectivity, large-scale networks and neuronal dynamics in schizophrenia. Current Opinion in Neurobiology, 23(2):283–290.

Uhlhaas, P. J., Linden, D. E., Singer, W., Haenschel, C., Lindner, M., Maurer, K., and Rodriguez, E. (2006). Dysfunctional long-range coordination of neural activity during Gestalt perception in schizophrenia. The Journal of neuroscience : the official journal of the Society for Neuroscience, 26:8168–75. 31.

Uhlhaas, P. J., Pipa, G., Lima, B., Melloni, L., Neuenschwander, S., Nikolic, D., and Singer, W. (2009). Neural synchrony in cortical networks: history, concept and current status. Front Integr Neurosci, 3:17.

Uhlhaas, P. J., Roux, F., Rodriguez, E., Rotarska-Jagiela, A., and Singer, W. (2010). Neural synchrony and the development of cortical networks. Trends in cognitive sciences, 14:72–80. 2.

Uhlhaas, P. J. and Singer, W. (2010). Abnormal neural oscillations and synchrony in schizophrenia. Nature reviews. Neuroscience, 11:100–13. 2.

Wendling, F., Bellanger, J. J., Bartolomei, F., and Chauvel, P. (2000). Relevance of nonlinear lumped-parameter models in the analysis of depth-EEG epileptic signals. Biological Cybernetics, 83(4):367– 378.

